# Distinct contributions of anterior and posterior orbitofrontal cortex to outcome-guided behavior

**DOI:** 10.1101/2025.10.08.681180

**Authors:** Qingfang Liu, Daria Porter, Hadeel Damra, Yao Zhao, Joel L. Voss, Geoffrey Schoenbaum, Thorsten Kahnt

## Abstract

The lateral orbitofrontal cortex (OFC) is critical for flexibly adjusting choices when outcome values change. Anterior and posterior parts of the human lateral OFC differ in cytoarchitecture and connectivity, but whether these subregions make differential contributions to outcome-guided (i.e., goal-directed) behavior remains unclear. Outcome-guided behavior requires (a) representations of stimulus–outcome associations and (b) inferring the current value of options when making decisions. Here, we test whether these two functions are differentially supported by the posterior (pOFC) and anterior (aOFC) parts of the lateral OFC, using transcranial magnetic stimulation (TMS) to selectively disrupt activity in functional networks centered on the pOFC and aOFC during a two-day outcome devaluation task. Participants (n = 48) received pOFC or aOFC network-targeted TMS either on day 1 before learning associations between visual stimuli and sweet or savory food odors, or on day 2 before a meal that selectively devalued one of these outcomes, followed by a choice test. TMS targeting pOFC, but not aOFC, before the meal on day 2 disrupted outcome-guided behavior, as measured by choices of stimuli predicting non-sated rewards in the post-meal choice test. In contrast, TMS targeting aOFC, but not pOFC, before learning on day 1 similarly impaired behavior in the post-meal choice test on day 2. These findings demonstrate that anterior and posterior parts of the lateral OFC make distinct contributions to outcome-guided behavior by supporting learning of stimulus–outcome associations and inferring the current value of options, respectively.

## INTRODUCTION

Humans and animals effortlessly adapt to changing environments by flexibly adjusting their behavior. This adaptability relies on outcome-guided (i.e., goal-directed) decision-making, where individuals re-evaluate their choices in real time, simulating potential outcomes based on changes in outcome value^1^ rather than defaulting to habitual responses. For example, a restaurant chef might anticipate that a guest could experience an allergic reaction to certain ingredients and adjust the dish accordingly before an issue arises. To enable this flexibility, a detailed representation of the environment—commonly referred to as a cognitive map or model—is essential.^2^ A chef with full knowledge of ingredients and associated allergies can efficiently modify recipes to accommodate allergies without compromising the dish.

The orbitofrontal cortex (OFC) has been proposed to play a central role in both processes, supporting adaptive behaviors through the formation of cognitive maps^3–5^ as well as their use to infer potential outcomes.^6, 7^ However, the OFC is a heterogeneous region, comprising multiple subregions with varying anatomical and functional properties along both medial-lateral and anterior-posterior axes.^8–17^ Whereas medial OFC has been linked to representing and comparing values at the time of choice,^18–20^ lateral OFC is implicated in representing, learning, and using stimulus–outcome associations, as revealed by studies on credit assignment,^21–23^ outcome-specific signaling,^19, 24–26^ model-based inference, and outcome devaluation.^6, 27–31^ Although the anterior-posterior axis has received comparatively less attention, functional imaging studies in humans suggest that posterior OFC responds to concrete, sensory-based outcome, whereas anterior OFC processes more abstract rewards.^32^ Moreover, in rodents, posterior OFC lesions disrupt reversal learning and devaluation, whereas anterior lesions have limited impact.^11^ Finally, inactivation of posterior OFC in monkeys impairs value updating, while inactivation of anterior OFC disrupts the ability to apply updated values during choice.^33^

The current study investigates if anterior and posterior subregions of the lateral OFC make distinct contributions to adaptive behavior in an outcome devaluation task.^4, 6, 29, 33–41^ Outcome devaluation assesses responses to predictive stimuli following the selective devaluation of their associated outcomes, thereby revealing the capacity to align choices with updated goals and contexts. In outcome-specific versions of this task, different stimuli are first associated with different but equally preferred rewards. Next, one of the outcomes is selectively devalued (for instance by feeding it to satiety), and then decisions between stimuli are assessed in a choice test.^33, 42^

Contemporary theories of OFC function propose that OFC is required for adaptive behavior in this task because it supports on-the-fly inferences about the current value of the choice options.^43, 44^ However, more recent work shows similar deficits in this task when OFC activity is disrupted during initial learning of stimulus-outcome associations,^3^ paralleling other tasks that require inference based on associative task structures (e.g., sensory preconditioning).^45^ This suggests that OFC is critically involved in forming the specific associations that link predictive stimuli to outcomes (i.e., the task model) during initial learning,^46^ in line with neural recoding studies showing such associative information is represented in the OFC.^24, 26, 47–50^ While it is possible that behavioral impairments following disruption of OFC activity at these different time points reflect the same functional deficit (i.e., loss of the task model), it is also possible that they reflect sparable functions (i.e., learning and use of the model), which are potentially supported by different OFC subregions. Here, we directly test these ideas by modulating the activity of two different OFC networks either before initial learning or before the choice test of the devaluation task.

Previous studies in non-human primates suggest that anterior and posterior regions of the OFC support distinct functions in outcome-guided behavior.^33^ Our earlier work further demonstrated that the posterior OFC in humans is critical for using stimulus–outcome associations in the devaluation task.^6^ Building on these findings, we hypothesized that the anterior and posterior subregions of lateral OFC support different functions required in the outcome devaluation task: the anterior OFC supports the acquisition of stimulus– outcome associations, and the posterior OFC supports their use in guiding choices. To test this, we applied network-targeted transcranial magnetic stimulation (TMS) either before initial training or before the choice test. This approach allowed us to test the specific roles of anterior and posterior OFC networks for learning associative structures and guiding choices based on current values.

Our findings reveal distinct roles for the anterior and posterior lateral OFC networks in outcome-guided behavior. Disruption of the posterior but not anterior lateral OFC network before the choice test impaired adaptive behavior, whereas disruption of the anterior but not posterior lateral OFC before initial learning similarly impaired subsequent outcome-guided behavior in the choice test. Together, these results suggest that anterior and posterior lateral OFC networks play complementary roles for outcome-guided behavior, supporting the acquisition and use of outcome-specific stimulus-reward associations, respectively.

## RESULTS

### Experimental design and outcome devaluation task

To separate the learning and use of stimulus-outcome associations, we utilized a two-day variant of an outcome devaluation task, in which learning of the specific stimulus-outcome pairings took place on Day 1, while selective devaluation took place on Day 2, bracketed by a pre- and post-meal choice test (Figure 1A).

**Figure 1.**
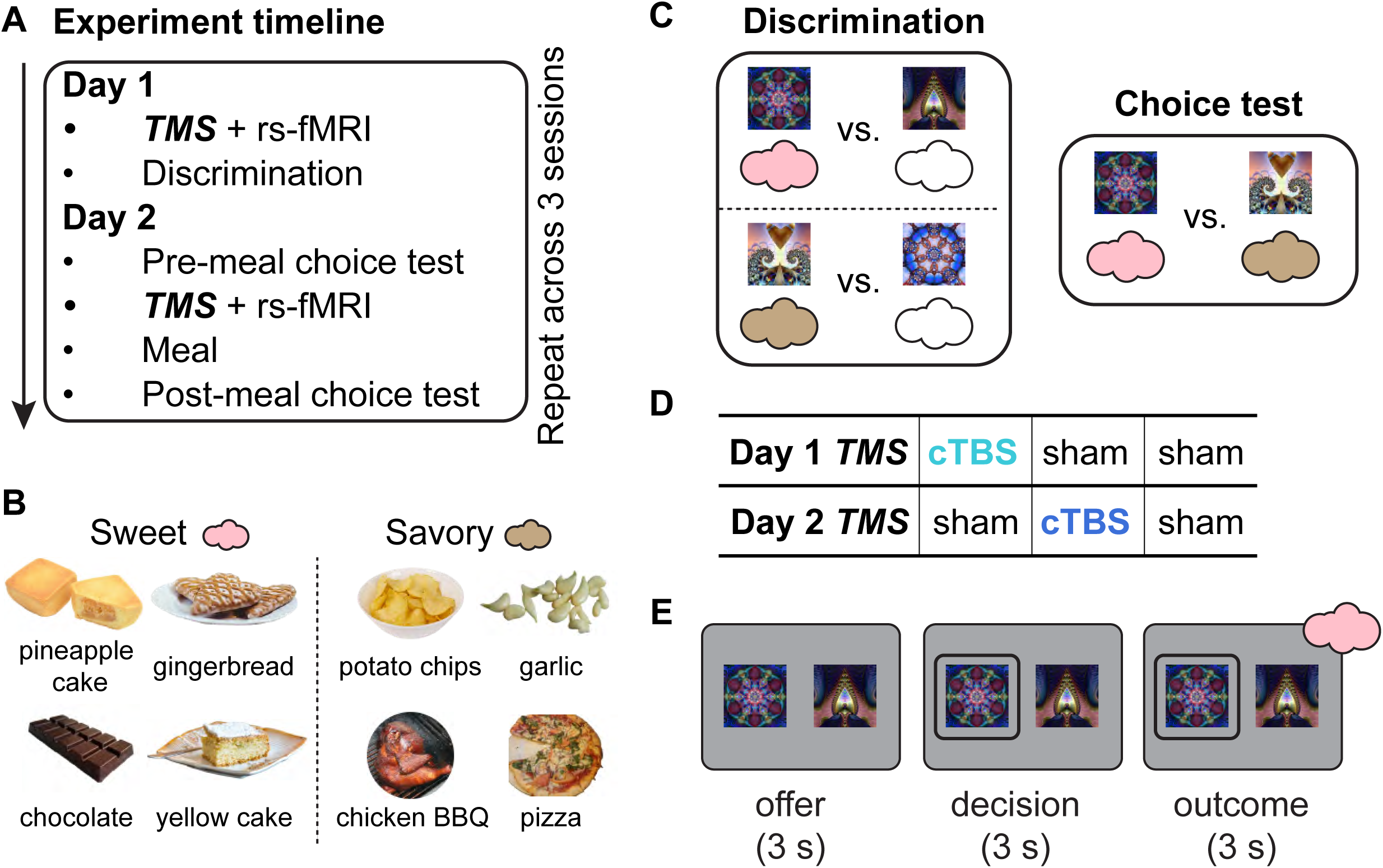
Experimental design and outcome devaluation task. **A.** Experiment timeline. On Day 1, participants received either continuous theta burst stimulation (cTBS) or sham TMS before a discrimination task. On Day 2, they performed a pre-meal choice test, received TMS (cTBS or sham), consumed a meal, and then completed a post-meal choice test. **B.** Odor stimuli. One savory and one sweet food odor (matched in pleasantness) was selected for each participant out of eight possible options. **C.** Task structure. In the discrimination task, participants learned which stimuli predicted odors (colored clouds) versus no odors (i.e., clean air, empty clouds). In the choice test, participants selected stimuli based on learned odor associations and their odor preference. **D.** TMS conditions across the three sessions. Each participant completed three two-day sessions, receiving cTBS either before learning (Day 1) or before the meal (Day 2), or sham on both days. This yielded three within-participant conditions: cTBS–sham, sham–cTBS, and sham–sham (order counterbalanced). The panel shows one example schedule. **E.** Trial structure of discrimination and choice tests. Each trial started with an offer phase (3 s), presenting two visual stimuli paired with different outcomes, followed by a decision phase (maximum 3 s) where participants selected one stimulus. In the discrimination task, the trial concluded with an outcome phase (3 s) where participants received an odor or no odor, depending on their choice.

On Day 1, participants learned to discriminate pairs of visual stimuli, where one stimulus was associated with a desirable food odor (sweet or savory, equally valued based on pre-task ratings; Figure 1B) and the other stimulus was associated with clean air (Figure 1C, left). Participants were asked to select the stimulus associated with any odor, meaning they were not incentivized to encode the specific stimulus-outcome identity association to perform the discrimination task.

On Day 2, participants made choices between cues predictive of different food odors both before and after a meal intended to selectively devalue one of the two food odors. In this choice test, participants made preference-based choices between stimuli predicting sweet and savory odors (Figure 1C, right). Participants received the odors associated with the chosen stimulus during the Day 1 discrimination task and the Day 2 pre-meal choice test, but no odors were delivered during the Day 2 post-meal choice test. Participants also reported how much they liked each odor before and after the meal.

Each participant repeated this two-day task three separate times (i.e., “sessions”; at least one week apart), each time learning new stimulus-food odor pairings. To test the role of OFC networks in learning and using stimulus-outcome associations, cTBS was administered at two different time points—either before the discrimination task on Day 1 or before the meal on Day 2 (Figure 1D). That is, each day, participants could receive either theta-burst stimulation (cTBS) or sham TMS, resulting in three within-participant conditions spread across the three sessions (Day 1–Day 2: cTBS–sham, sham–cTBS, sham– sham; order counterbalanced across participants; Figure 1D). When applied over motor cortex, this cTBS protocol (cTBS_600_) reduces cortical excitability for about 50 minutes.^51^

### TMS targeting dissociable anterior and posterior OFC networks

To test the potentially distinct functional roles of OFC subregions in learning and using stimulus-outcome associations, TMS targeted either the anterior (aOFC) or posterior (pOFC) portions of the lateral OFC in different groups of participants (N=23 and N=25; Figure 2A). Stimulation targets were defined on resting-state fMRI data collected on an initial study visit (before the three experimental sessions), by seeding a functional connectivity analysis in the right hemisphere: aOFC at MNI coordinates [34, 54, −14] and pOFC at [28, 38, −16]. We individually identified stimulation sites in lateral prefrontal cortex (LPFC) ROIs (referred to as aOFC-conn-LPFC and pOFC-conn-LPFC, respectively) that exhibited the highest functional connectivity with either the aOFC or pOFC seed region (Figure 2B).

**Figure 2:**
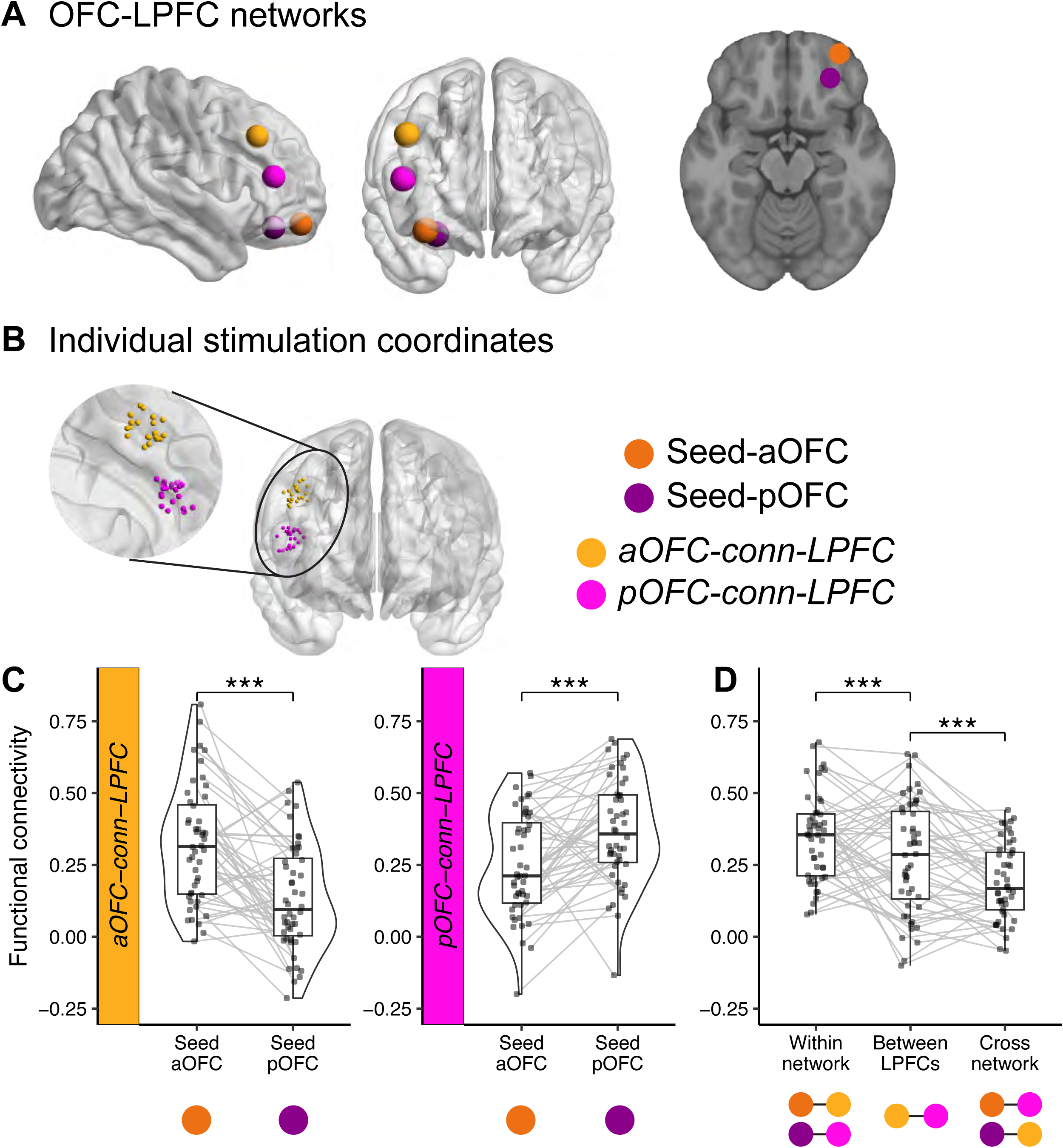
TMS targeting dissociable anterior and posterior OFC networks. **A.** OFC-LPFC networks. Seed regions in the anterior (aOFC; tangerine, MNI coordinates: [34, 54, –14]) and posterior OFC (pOFC; magenta, MNI coordinates: [28, 38, –16]), along with their corresponding connectivity-based target regions in the lateral prefrontal cortex (LPFC), are shown on cortical surface renderings. Brain visualizations were generated using BrainNet Viewer,^78^ and the axial slice corresponds to *z* = −16 in MNI space. **B.** Individual stimulation coordinates. LPFC stimulation sites were individually selected to maximize functional connectivity with either the aOFC or pOFC seed region. The zoomed view shows the distribution of stimulation coordinates across participants, color-coded by group. **C.** Functional connectivity estimates. Half-violin plots depict the distribution of resting-state functional connectivity between LPFC stimulation sites and each OFC seed region. Each dot represents an individual participant’s connectivity estimate, and gray lines connect seed regions from the same subject. Boxplots indicate the median and interquartile range. Asterisks denote significant differences between connectivity patterns (***p *<* 0.001). **D.** Functional connectivity within-network, between-LPFC, and cross-network. Boxplots show functional connectivity across participants for within-network (OFC seed to its corresponding LPFC target), between-LPFC (connectivity between the two LPFC targets), and cross-network (OFC seed to the non-corresponding LPFC target) connections. Asterisks denote statistical significance (***p *<* 0.001).

Resting-state fMRI data were collected immediately after administering TMS on each Day 1 and Day 2 visit. We confirmed the functional dissociation between anterior and posterior OFC networks across all resting-state fMRI scans: the aOFC-conn-LPFC site showed significantly stronger connectivity with the aOFC seed than with the pOFC seed (*p <* 2.2 × 10^−16^, linear mixed-effects model), and the pOFC-conn-LPFC site showed significantly stronger connectivity with the pOFC seed than with the aOFC seed (*p <* 2.2 × 10^−16^, linear mixed-effects model) (Figure 2C).

To further assess the specificity of these networks, we compared within-network, between-LPFC, and cross-network functional connectivity (Figure 2D). Within-network connectivity was defined as the average connectivity between each OFC seed (aOFC or pOFC) and its corresponding LPFC target, whereas cross-network connectivity was defined as the connectivity between each OFC seed and the non-corresponding LPFC target. Between-LPFC connectivity was measured as the functional connectivity between the two LPFC targets. Across participants, within-network connectivity was significantly stronger than between-LPFC connectivity (*p* = 7.495 × 10^−9^, linear mixed-effects model), and between-LPFC connectivity was significantly stronger than cross-network connectivity (*p* = 1.503 × 10^−12^). Together, these findings demonstrate that our TMS approach targeted dissociable OFC networks.

### Discrimination learning and selective satiation effects

Over the five blocks of the discrimination task on Day 1, choices of odor-predictive stimuli (vs. clean air) increased significantly across blocks in both aOFC and pOFC groups, indicating successful discrimination learning (Figure 3A; *p <* 2.2 × 10^−16^, linear mixed-effect models). Although this increase was influenced by TMS applied prior to the discrimination task (cTBS vs. sham; *p* = 1.24 × 10^−7^), there was no effect of cTBS on choices of odor-predicting stimuli in the last block of the discrimination task in either group (aOFC: *p* = 0.605; pOFC: *p* = 0.967, t-test). There was also a significant main effect of session number (Figure S1A; 1^st^, 2^nd^, 3^rd^; *p* = 1.66 × 10^−11^) as well as a significant session-by-TMS interaction (Figure S1B; *p* = 1.90 × 10^−5^).

**Figure 3:**
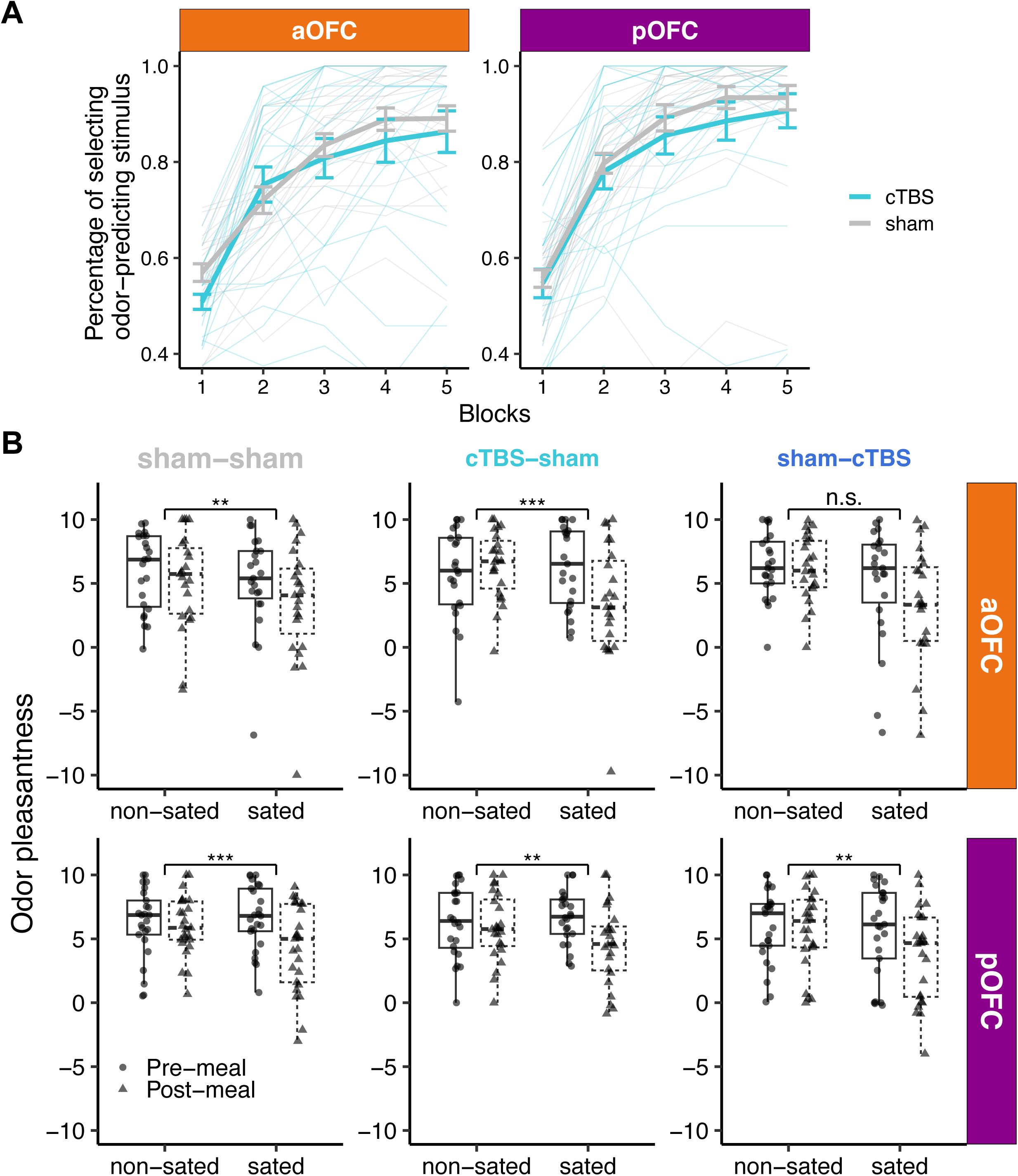
Discrimination learning and selective satiation effects. **A**. On day 1, participants in both the aOFC and pOFC groups learned to select odor-predictive stimuli across five blocks of the discrimination task. Thin lines show individual trajectories; thick lines with error bars indicate group means ± s.e.m. for cTBS and sham. **B**. Pre- and post-meal odor pleasantness ratings separated by TMS condition (sham–sham, cTBS–sham, sham–cTBS), stimulation target (aOFC vs. pOFC), and odor type (sated vs. non-sated odor). These ratings confirm that selective satiation effects were evident across most TMS conditions within each group. Asterisks denote statistically significant time-by-odor interaction (∗∗p < 0.01, ∗∗∗p *<* 0.001, n.s. = not significant). *Data plotted are on the original rating scale; however, statistical analyses were conducted on normalized ratings to account for individual differences in scale use.* See also Figure S1 for additional details on discrimination learning and Figure S3 for correlations between selective satiation effects and overall probe choices.

On Day 2, participants were given the opportunity to eat a meal that was matched to either the sweet or the savory food odor. To evaluate the effectiveness of the meal in selectively devaluing the meal-matched odor, we examined changes in odor pleasantness ratings from before to after the meal. Selective satiation robustly reduced the rated pleasantness of the meal-matched odor compared to the non-matched odor (post-meal minus pre-meal) (*p* = 2.75 × 10^−13^; Figure 3B), regardless of TMS condition (sham vs. cTBS, Day 2), stimulation target (aOFC vs. pOFC), session number (1^st^, 2^nd^, 3^rd^), or sated odor type (savory/sweet) (all *p >* 0.05). Consistent with previous findings,^6, 52, 53^ these results demonstrate that disrupting OFC activity does not affect the ability to devalue rewards themselves.

Taken together, these results show that participants in all groups and TMS conditions learned the discrimination on Day 1 and that the meal on Day 2 selectively devalued the meal-matched odor.

### Posterior OFC-targeted cTBS before the meal impairs outcome-guided behavior

Before and after the meal on Day 2, participants performed a choice test where they chose between stimuli predicting the sated and non-sated odor. Across groups and conditions, participants’ choices showed a significant effect of time (pre- vs post-meal), with a significant decrease in choices of sated odor-predicting stimuli from pre- to post-meal (Wilcoxon signed-rank test, two-sided, *V* = 3062.5, *p* = 5.31 × 10^−4^).

To examine the contribution of the aOFC and pOFC to using the stimulus-outcome associations learned on Day 1 to flexibly infer the current value of the choice options, we compared choices of sated odor-predicting stimuli in this choice test between conditions where cTBS or sham TMS was applied before the meal on Day 2 (i.e., “sham-sham” and “sham-cTBS”). Because post-meal choices were significantly driven by a number of factors (pre-meal choices, selective satiation, learned stimulus value on Day 1 (*w_SA_* − *w_NS_*), see Figure S2), we modeled post-meal choices using logistic mixed-effects models (see Methods), accounting for these factors (Figure 4B; see Figure S5A for raw data without accounting for covariates). In the pOFC group, we found that cTBS compared to sham significantly increased choices of sated odor-predicting stimuli (*p* = 0.00036), indicating poorer adaptation to the updated value of the outcomes. No effect of cTBS was found in the aOFC group (*p* = 0.655), and the difference between the aOFC and pOFC group was significant, as indicated by a significant group-by-TMS (sham vs. cTBS on Day 2, Day 1 fixed at sham) interaction (*p* = 0.00565).

**Figure 4:**
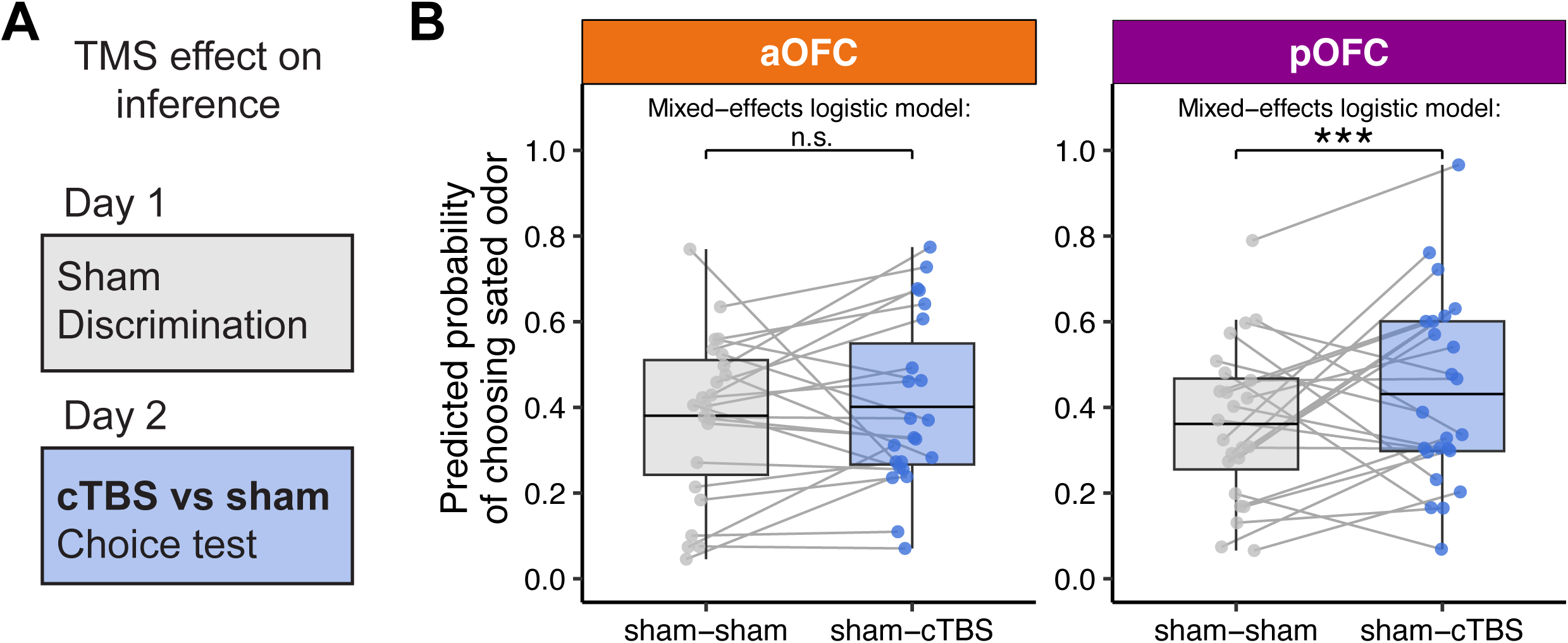
Posterior OFC-targeted cTBS before the meal impairs outcome-guided behavior. **A**. To test the role of OFC networks in using stimulus-outcome associations to infer the current value of choice options, we compared post-meal choices on Day 2 between conditions when cTBS or sham TMS was applied before the meal on Day 2. **B**. Predicted probability of choosing sated odor-predicting stimuli in sham–sham and sham–cTBS conditions, shown separately for anterior (aOFC, tangerine) and posterior (pOFC, magenta) OFC-targeting groups. Each dot represents a participant’s average predicted probability, and gray lines connect values from the same participant across conditions. Box plots show group-level distributions of fitted values, with horizontal lines representing the group means. Statistical comparisons were conducted using trial-wise mixed-effects logistic regression controlling for baseline odor preference, satiation status, and value difference between sated and non-sated options (*w_SA_* − *w_NS_*). A significant increase in sated odor choice was observed following Day 2 pOFC cTBS (***p *<* 0.001), but not in the aOFC group (n.s.). See also Figures S2–S3 for covariates (learned stimulus values and satiation effects) accounted for by the results presented here; Figure S4A,B for model fit evaluation; Figure S5A for raw data without covariate adjustment; and Figure S6A,C for correlations of perceived TMS discomfort and intensity with probe choices.

To evaluate the fit of the mixed-effects models, we assessed for each group and condition, (1) the correlation between each subject’s mean predicted probability of choosing the sated odor-predicting stimulus and their actual mean choice rate, as well as (2) the model’s trial-level discrimination ability via ROC analysis (Figure S4AB). The high between-participant correlation indicate that the model effectively captured individual differences in choice behavior. ROC curves further demonstrated reliable trial-level discrimination, with AUCs ranging from 0.71 to 0.78 across conditions and stimulation targeting groups.

We conducted additional analyses to assess whether the effect of TMS on sated odor-predicting stimulus choices was driven by other factors, such as satiation status or perceived TMS discomfort or intensity. The between-participant correlations between selective pleasantness changes and post-meal choices were not affected by Day 2 cTBS (all *p >* 0.05; Figure S3A), suggesting that the effect of Day 2 cTBS on choices was not modulated by selective satiation. Moreover, TMS-related changes in post-meal choices could not be explained by perceived TMS discomfort or intensity, as incorporating TMS ratings into the regression models did not alter any of the findings (Figure S6).

Together, these results show that pOFC-targeted cTBS before the meal impairs outcome-guided behavior, as indicated by the continued selection of stimuli that predict sated odors. In contrast, aOFC-targeted cTBS had no such effect, and suggesting that pOFC but not aOFC plays a critical role in using stimulus-outcome associations to infer the current value of choice options.

### Anterior OFC targeted cTBS before discrimination learning impairs outcome-guided behavior

The previous results show that disrupting pOFC network activity impairs behavior in the post-meal choice test, suggesting a role of pOFC in inferring the current value of the choice options based on stimulus-outcome associations learned on Day 1. This deficit could be either due to a disruption of the inference process as such, or a disruption of encoded stimulus-outcome associations that are required for this inference. If the latter is true, modulating OFC network function during discrimination learning on Day 1 should have similar effects on post-meal choices on Day 2. Alternatively, it is possible that learning and use of stimulus-outcome associations are spatially dissociable within the primate OFC, potentially along its anterior-posterior axis. To test these questions, we examined post-meal choices on Day 2 in sessions where cTBS or sham TMS targeting the aOFC and pOFC network was applied before the discrimination task on Day 1 (i.e., “sham-sham” and “cTBS-sham” conditions, Figure 5A).

**Figure 5:**
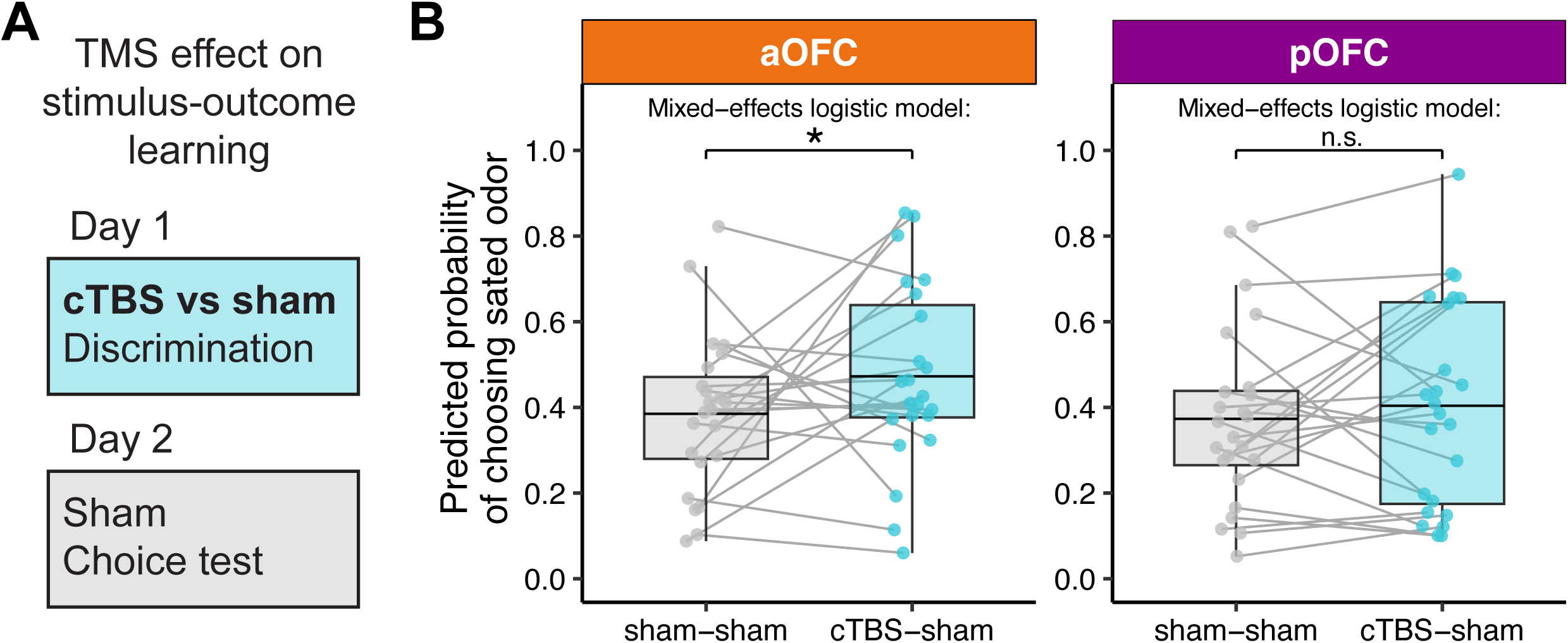
Anterior OFC-targeted cTBS before discrimination learning impairs outcome-guided behavior. **A**. To test the role of OFC networks in learning specific stimulus-outcome associations, we compared post-meal choices on Day 2 between conditions when cTBS or sham TMS was applied before the discrimination task on Day 1. **B**. Predicted probability of choosing the sated odor in the post-meal test, compared between sham–sham and cTBS–sham sessions, separately for anterior (aOFC, tangerine) and posterior (pOFC, magenta) targeting groups. Each dot represents a participant’s average probability of choosing the sated odor-predicting stimulus as predicted by the mixed-effects model, with gray lines connecting values from the same participant across conditions. Box plots show group-level distributions of fitted values, with horizontal lines representing the group means. Statistical comparisons were conducted using trial-wise logistic mixed-effects models, controlling for value difference, pre-meal choices, and selective satiation. A significant increase in sated odor choice was observed following Day 1 cTBS compared to sham in the aOFC group (p *<* 0.05*), but not in the pOFC group (n.s.). See also Figures S2– S3 for covariates (learned stimulus values and satiation effects) accounted for by the results presented here; Figure S4C,D for model fit evaluation; Figure S5B for raw data without covariate adjustment; and Figure S6B,D for correlations of perceived TMS discomfort and intensity with probe choices.

As above, because choices of sated odor-predicting stimuli in the post-meal choice test were significantly influenced by a number of factors (pre-meal choices, selective satiation, learned stimulus value on Day 1 (*w_SA_*− *w_NS_*)), we modeled post-meal choices using logistic fixed-effects models (Figure 5), accounting for these factors (see Figure S5 for raw data without accounting for covariates). In the aOFC group, we found that compared to sham, cTBS on Day 1 significantly increased choices of sated odor-predicting stimuli in the post-meal choice test on Day 2 (Figure 5B; *p* = 0.015). There were also a significant effect of session number (*p* = 8.5 × 10^−5^) and a significant TMS-by-session interaction (*p* = 0.024), indicating that the effect of cTBS diminished over sessions.

In contrast, similar analyses in the pOFC group revealed no significant effect of cTBS applied before the discrimination task on Day 1on post-meal choices on Day 2 (Figure 5B; *p* = 0.24). However, no significant interaction between group (aOFC vs. pOFC) and TMS condition (sham-sham vs. cTBS-sham) was found (*p* = 0.37). Again, these analyses accounted for various factors, as pre-meal choices and value difference (but not selective satiation) were significant predictors of post-meal choices. Model fits were evaluated using the same approach as for testing effects of TMS on Day 2, combining across-participant correlation and ROC analysis to assess prediction accuracy at both the participant and trial levels (Figure S4CD).

Taken together, these results show that aOFC-targeted cTBS before the conditioning on Day 1 impairs outcome-guided behavior on Day 2, as indicated by the continued selection of stimuli that predict sated odors. In contrast, pOFC-targeted cTBS had no such effect, suggesting that aOFC but not pOFC plays a critical role in learning stimulus-outcome associations during the discrimination task on Day 1.

### cTBS distorts low-dimensional connectivity structures

In a final step, we tested the effects of TMS on the resting-state fMRI data collected following all TMS applications (Figure 1), along with a baseline (i.e., null) scan on the initial study visit. To identify neural evidence of cTBS-induced effects on OFC networks, we applied a conditional variational autoencoder (cVAE) to these resting-state fMRI data.^54–57^ A variational autoencoder (VAE) is an unsupervised deep generative model that learns low-dimensional latent representations from high-dimensional inputs. We used a modified version—cVAE conditioned on participant identity—which allows the model to account for individual differences in functional connectivity profiles while capturing stimulation-related differences in the latent space (Figure 6A).

**Figure 6:**
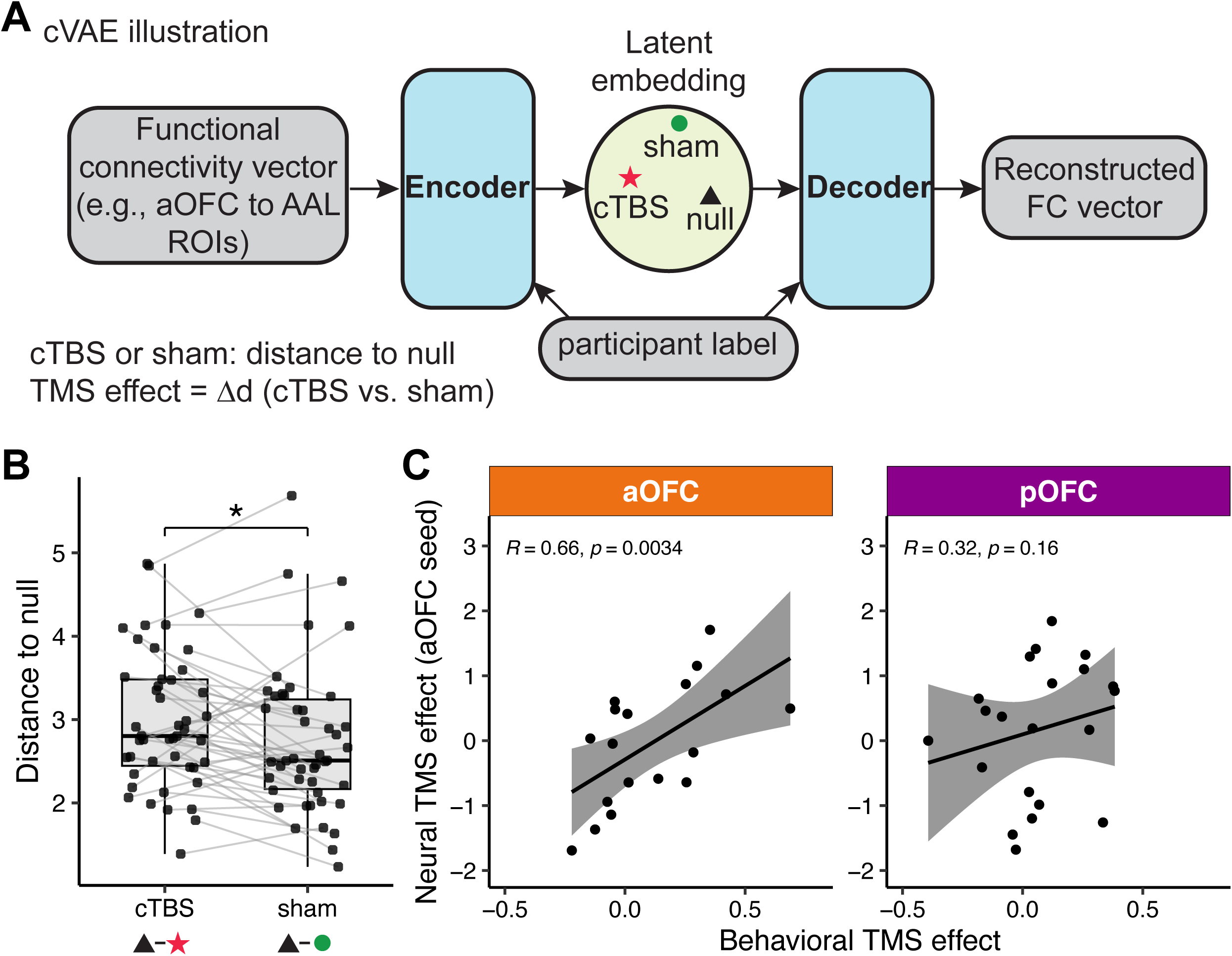
cTBS distorts low-dimensional connectivity structures. **A**. Illustration of the conditional variational autoencoder (cVAE) used to encode functional connectivity patterns (e.g., between aOFC and AAL ROIs) into a latent space, conditioned on participant. We tested connectivity patterns for each target and seed ROI separately. The neural network reconstructs functional connectivity patterns via encoding and decoding layers and computes the distance from each condition (cTBS or sham) to a reference null distribution. The neural TMS effect is defined as the distance between cTBS and sham conditions. **B**. Paired distances to the null in the latent space for cTBS vs. sham conditions using functional connectivity patterns between aOFC and AAL ROIs. Most participants showed greater distances under cTBS, confirmed by a paired t-test (*t*(45) = 2.67, *p* = 0.011). **C**. Correlation between the effect of cTBS on post-meal choices (Day 1 comparison, sham–sham vs. cTBS–sham) and neural cTBS effects from panel B. In the aOFC group, neural TMS effects are associated with larger effects on post-meal choices of sated odor predicting stimuli (*R* = 0.54, *p* = 0.027), while no such relationship is observed in the pOFC group (*R* = 0.31, *p* = 0.18).

We treated the initial resting-state scan as a reference and hypothesized that cTBS would induce greater deviation from this baseline compared to sham. Thus, for each resting-state scan, we computed the Euclidean distance to this baseline in the latent space learned by the cVAE. To identify potential cTBS effects, we extracted the pattern of functional connectivity between each seed or stimulation ROI (Figure 2A) and a set of Automated Anatomical Labeling (AAL) ROIs and analyzed them separately. Using functional connectivity patterns between the aOFC seed and AAL ROIs, we found that cTBS was associated with a significantly greater deviation from baseline than sham TMS (t(45) = 2.41, p = 0.020), indicating a reliable modulation effect in the aOFC (Figure 6B). No such effect was observed when using other seed or stimulation ROIs.

We further tested whether the condition-wise neural TMS effect derived from the aOFC seed ROI could account for individual differences in behavior (Figure 6C). In the aOFC group, TMS effects on connectivity were significantly correlated with the behavioral effects of cTBS on Day 1 on post-meal choices on Day 2 (R = 0.66, p = 0.0034), whereas no significant relationship was observed in the pOFC group (R = 0.32, p = 0.16). This finding further supports the idea that anterior, but not posterior, OFC contributed to learning of specific stimulus–outcome associations on Day 1 (Figure 5).

## DISCUSSION

In this study, we used network-targeted TMS in the context of an outcome devaluation task to selectively modulate activity in networks centered on anterior and posterior subregions of the human lateral OFC either during learning of stimulus-outcome associations or prior to a meal and subsequent choice test. We found that TMS targeting the posterior OFC network prior to the meal disrupted outcome-guided behavior, as evidenced by continued choices of stimuli predicting sated rewards in the post-meal choice test. Conversely, targeting the anterior OFC network before learning stimulus-outcome associations also impaired behavior in the post-meal choice test.

These findings suggest that the OFC makes a two-fold contribution to outcome-guided behavior. First, it supports the acquisition of stimulus-outcome associations, that is, the construction a model of the associative task structure. Second, it is involved in using this model to infer the current value of the choice options. Importantly, our results suggest that these functions are carried out by different subregions of OFC. Whereas anterior OFC contributes to model construction, posterior OFC contributes to using this model for inference.

The involvement of OFC in model construction and use is in line with previous correlational and causal work across species. For instance, OFC activity correlates with specific stimulus-outcome and stimulus-stimulus associations in rats,^26, 50, 58^ non-human primates,^59^ and humans,^24, 25, 47, 49^ and disruption of OFC activity using lesions, optogenetics, pharmacological inactivation or network-targeted TMS after learning is completed causes deficits in using these associations to guide behavior.^6, 27, 31, 42, 60–63^ Moreover, work in rats shows that inactivation during initial learning of the task structure similarly impairs the later use of the model at a time when OFC is undisturbed.^3, 45^ The finding that modulation of OFC network activity during both learning and choice impairs outcome-guided behavior could be taken to suggest that the key contribution of OFC is to learn and store the model.

However, our current results suggest that model construction and use are separate functions that are supported by distinct parts of lateral OFC. Specifically, our findings suggest a segregation of function within lateral OFC, such that anterior OFC constructs the model and posterior OFC uses the model for inferring the current value of choice options. This contrasts with work in rats, where inactivation of the same parts of OFC during learning and choice cause similar deficits in outcome-guided behavior. This discrepancy between findings in rats and humans may be attributable to species differences in OFC anatomy. The primate OFC consists of both agranular and granular cortex, but the rat OFC is exclusively agranular.^64^ Intriguingly, the posterior OFC in primates is predominantly agranular, whereas the anterior OFC is granular.^65^ Moreover, primate anterior and posterior OFC also differ in terms of connectivity,^10, 66, 67^ such that posterior OFC receives input from sensory cortices and anterior OFC receives inputs from association cortices.^65^ Of note, anterior and posterior subregions of the lateral OFC of rats also differ in connectivity,^68^ and it is possible that the functional differentiation shown here is also present in rats but has been overlooked due to its small size.

Our findings suggest that the anterior OFC plays a critical role in learning specific stimulus-outcome associations even when the task does not explicitly require it. That is, although our discrimination task involved rewarding outcomes, learning the specific identity of rewards was not reinforced or required for performance. Such latent learning parallels previous research indicating that both humans and animals construct a representation of the task environment even in the absence of rewards.^5, 69–71^ This information, once formed, is the foundation for outcome-guided behaviors.^2, 69^ As such, the effect of disrupting this latent learning can be revealed in later stages, when the information becomes crucial for behavior.

Replicating our previous work,^6^ targeting pOFC disrupted post-meal choices, suggesting that pOFC contributes to updating the current value of choice options. At its face, an alternative explanation of this finding is that pOFC is required for value comparison and choice per se. However, the former interpretation is supported by work in animals indicating that OFC is not required for value-based choice as such, but only for updating the value of reward-associated stimuli following devaluation.^27, 28, 63^ While our data are consistent with this account, alternative explanations remain possible.

The functional heterogeneity of OFC revealed here aligns with and extends prior studies demonstrating distinct roles of OFC subregions across various tasks and species, including outcome devaluation,^33^ two-choice probabilistic tasks,^59^ encoding of value information,^72^ and economic decision-making.^12^ Particularly relevant is work in non-human primates showing differential roles of OFC subregions in outcome devaluation, with anterior OFC (area 11) being involved in goal selection during choice and posterior OFC (area 13) supporting value updating.^33^ In contrast to this work, our study focused on the differential involvement of lateral OFC subregions in learning and using stimulus-outcome associations. Identifying such functional differences within OFC advances our understanding of how this large and heterogeneous brain region supports learning and behavior.

Our TMS approach aimed to target specific networks, rather than isolated areas. That is, the LPFC stimulation sites were selected because they are part of the same functional network as the anterior or posterior OFC targets. As such, we interpret the behavioral effects of TMS as being driven by neural changes in both areas, and/or functional interactions between them. However, our rs-fMRI data analysis revealed a correlation between the TMS effects on behavior and fMRI connectivity only in the targeted aOFC area, suggesting the behavioral effects are mediated by changes in the targeted area (at least in aOFC). Similarly, our previous study^6^ found a relationship between effects of TMS on behavior and connectivity in the targeted OFC but not in the directly stimulated LPFC. In this context, it is important to note that we only found an effect of TMS on connectivity in the aOFC but not in the pOFC. This may be due to insufficient power or other factors, given that our previously^6, 49^ found effects of pOFC-targeted TMS on pOFC connectivity using different experimental designs, tasks, and sample sizes. Moreover, we did not find a significant interaction between the aOFC and pOFC groups, leaving open the possibility that pOFC-targeted stimulation also modulated aOFC activity to some extent. However, these unspecific effects in the pOFC group were not related to task behavior.

Although unexpected, we found that cTBS targeting both the anterior and posterior OFC disrupted discrimination learning on Day 1, especially during the first session. This challenges the view that OFC plays no role in simple Pavlovian learning,^26, 73, 74^ in line with recent rodent work suggesting that OFC’s role in Pavlovian acquisition may be more nuanced than previously thought.^75^ It is also in line with previous work showing that OFC supports learning in tasks that involve different reward identities, such as here.^76^ However, this deficit emerged only in the first session in our experiment, suggesting it may reflect an impairment in understanding the basic task structure. Once the structure was learned, it could be reused in subsequent sessions with different stimulus sets.^2, 77^ To account for these effects, we included the stimulus-level learned values of each option in the analysis of post-meal choices, rather than assuming equal learning.^6, 33^

In this regard, our within-participant design is a limitation, as it complicates interpretation. For instance, participants could learn during the first session that odor identity would be important on Day 2, potentially altering their strategies in later sessions. To test this possibility, we compared groups of participants based on the order of cTBS and sham stimulation. Although our findings were not affected by session order, the small sample size within each session-order group may have limited our ability to detect subtle order effects. Another limitation is the difference in perceived TMS intensity and discomfort between cTBS and sham conditions as reported in the current study and our previous work.^49^ However, we found no differences in these ratings between groups receiving TMS targeting the anterior and posterior OFC, and individual differences did not account for the observed behavioral effects.

In conclusion, our findings reveal distinct roles of anterior and posterior OFC in the formation and use of cognitive maps for outcome-guided behavior, advancing our understanding of how the OFC contributes to outcome-guided behavior.

## RESOURCE AVAILABILITY

### Lead contact

Further information and requests for resources should be directed to and will be fulfilled by the lead contact, Thorsten Kahnt.

### Materials availability

This study did not generate new unique materials or reagents.

### Data and code availability

#### Data

- The NODEAP fMRI dataset generated and analyzed during this study has been deposited at OpenNeuro and is publicly available as of the date of publication.
- DOI: https://openneuro.org/datasets/ds006693/versions/1.2.0
- The dataset is shared under a CC0 license and includes all raw and minimally processed BIDS-formatted MRI data, behavioral data, and relevant metadata.

#### Code

• All original preprocessing and analysis code has been deposited on GitHub and archived on Zenodo; it is publicly available at the following repositories:

– Main project code and processed behavioral data: https://github.com/QingfangLiu/project-nodeap-core
DOI: <u>10.5281/zenodo.17236831</u>

– cVAE analysis code and session-wise processed connectivity data from resting- state fMRI: https://github.com/QingfangLiu/project-nodeap-fmri-cvae
DOI: <u>10.5281/zenodo.17236867</u>
• No proprietary code was used in this study.

#### Additional Information

- Any additional information required to reanalyze the data reported in this paper is available from the lead contact upon request.

## ACKNOWLEDGMENTS

This work was supported by National Institute on Deafness and Other Communication Disorders grant R01DC015426 (to T.K.) and the Intramural Research Program at the National Institute on Drug Abuse (ZIA DA000642 to T.K. and DA000587 to G.S.). This research was supported in part by the Intramural Research Program of the National Institutes of Health (NIH). The contributions of the NIH authors are considered Works of the United States Government. The findings and conclusions presented in this paper are those of the authors and do not necessarily reflect the views of the NIH or the U.S. Department of Health and Human Services.

## AUTHOR CONTRIBUTIONS

Conceptualization, Q.L., T.K., and G.S.; Methodology, Q.L., T.K., G.S., and J.L.V.; Investigation, D.P., H.D., and Y.Z.; Formal analysis, Q.L., G.S., and T.K.; Writing – original draft, Q.L. and T.K.; Writing – review & editing, all authors; Supervision, T.K.

## DECLARATION OF GENERATIVE AI AND AI-ASSISTED TECHNOLOGIES

During the preparation of this work the authors used ChatGPT by OpenAI in order to check grammar. After using this tool, the authors reviewed and edited the content as needed and take full responsibility for the content of the publication.

## STAR⍰METHODS

### EXPERIMENTAL MODEL AND STUDY PARTICIPANT DETAILS

Eighty-eight healthy, right-handed participants (ages 18-40) with no history of psychiatric or neurological disease provided written informed consent to participate in this study. Of these, 48 participants (16 males; ages 18-40, mean = 25.17, SD = 4.14) completed all sessions. Due to a technical error, behavioral data from the cTBS-sham session were unavailable for one participant in the posterior targeting group; however, data from the other two sessions were included in the analysis where applicable. MRI data for five resting-state scans were not acquired and excluded from analysis. All participants fasted for at least four hours before each study visit.

### METHOD DETAILS

#### Study design

The study consisted of eight visits (Figure 1A, D), with Day 1 and Day 2 occurring on consecutive days. The two-day experiment was repeated across three sessions. Sessions were spaced at least one week apart, with a median interval of 13.5 days, a mean of 18.02 days (SD = 9.09), and a range of 7 to 63 days. On each Day 1 and Day 2, participants received either continuous theta-burst stimulation (cTBS, labeled C) or sham stimulation (S). Over the three sessions, they experienced three TMS conditions: cTBS-sham (CS), sham-cTBS (SC), and sham-sham (SS). The order of these conditions was counterbalanced, with 9 participants receiving CS-SC-SS, 7 receiving CS-SS-SC, and the remaining 32 equally assigned to one of the other four possible sequences (SC-CS-SS, SC-SS-CS, SS-CS-SC, and SS-SC-CS).

To prevent differences in stimulation location from affecting participants’ experience across sessions, each participant received TMS targeting either the anterior or posterior portion of the lateral OFC throughout all three sessions. Among the participants, 16 of 32 females and 9 of 16 males received TMS targeted to the posterior portion. Additionally, the order of satiation conditions was counterbalanced: half of the participants received a sweet meal in their first session, while the other half received a savory meal. The sated odor type alternated for each participant across the three sessions (e.g., savory-sweet-savory or sweet-savory-sweet).

#### Screening session

After providing informed consent and completing eligibility screening, participants rated the pleasantness of eight food odors. These odors, supplied by International Flavors and Fragrances (New York, NY), included four savory (garlic, potato chip, pizza, barbecue) and four sweet (chocolate, yellow cake, pineapple cake, gingerbread) odors. In each trial, participants smelled a food odor for 2 seconds and rated their liking on a visual analog scale ranging from “Most Disliked Sensation Imaginable” to “Most Liked Sensation Imaginable.” Ratings were made using a scroll wheel and keyboard press. Each odor was presented three times in a pseudo-randomized order, and ratings were averaged per odor. Based on these ratings, two odors (one savory, one sweet) that were pleasant (above neutral) and closely matched were selected for the discrimination and choice tests. These odors were used across all three sessions. Participants were excluded if no suitable odors were identified.

A custom-built, computer-controlled olfactometer was used to deliver the odors with precise timing to nasal masks worn by participants. The olfactometer directed medical-grade air through the headspace of amber bottles containing the odor solutions at a constant flow rate of 3.2L/min. Using two independent mass flow controllers (Alicat, Tucson, AZ), the device enabled precise dilution of the odorized air with odorless air. Throughout the experiment, a constant stream of odorless air was delivered, and odorized air was mixed in at specific time points without altering the overall flow rate or causing somatosensory stimulation.

#### Initial study visit: Scan & motor threshold

We acquired a T1-weighted structural MRI scan to assist with TMS neuronavigation and an 8 min multiecho resting-state fMRI scan (310 volumes, TR = 1.5s) to individually define the OFC-targeted cTBS coordinates. The same scanning parameters were used for all resting-state scans. We then measured resting motor threshold (rMT) by administering single TMS pulses to the hand area of the left motor cortex. rMT was defined as the lowest stimulator output required to evoke 5 visible thumb movements from 10 pulses.

#### Day 1: Discrimination task

Participants first underwent a TMS session (cTBS or sham) followed by a resting-state scan. They then completed five blocks of a discrimination task. In each trial, participants chose between two fractal stimuli: one associated with a savory or sweet odor, and the other with clean air. Stimuli were displayed for 3 seconds, followed by a choice phase (maximum 3 seconds). If participants selected a stimulus leading to an odor, the odor was delivered for 2 seconds. The inter-trial interval ranged from 4 to 8 seconds. Each run consisted of 24 trials, using four groups of stimulus pairs: two sets (A and B) crossed with sweet/savory odors. Each combination had three non-overlapping stimulus pairs, resulting in 24 distinct fractals. Each pair was presented twice to counterbalance left and right positions on the screen. Choice and response times were recorded for each trial, and different fractals were used across the three sessions.

#### Day 2: Meal consumption and choice test

Day 2 started with an odor pleasantness rating, followed by a pre-meal choice test where participants selected between pairs of stimuli. The pleasantness ratings were collected only for the two odors selected for each participant during the screening session and were measured three times. Afterwards, they underwent a TMS session and then had a meal carefully matched in flavor to either the sweet or savory food odor used in the task. Following the meal, participants completed another set of odor pleasantness ratings and a post-meal choice test. In both pre-meal and post-meal choice tests, participants were instructed to choose based on their current odor preferences.

The purpose of the meal was to selectively satiate one of the two food odors. Meal items were carefully chosen to closely match the corresponding food odors, and water was provided. Participants were given 15 minutes and instructed to eat until they felt very full. On average, participants consumed 669.89 ± 44.16 calories (SEM). Before analyzing the relationship between odor ratings and task behavior, ratings were standardized within each participant across sessions.

The pre-meal choice test included 30 trials, all from set A, consisting of 3 sweet vs. clean air pairs, 3 savory vs. clean air pairs, and 9 savory vs. sweet pairs. Each pair was presented twice to counterbalance left and right positions on the screen. The post-meal choice test included 60 trials from both sets A and B. In both pre- and post-meal choice tests, similar to the discrimination task, every trial began with a pair of stimuli presented for 3 seconds, followed by a decision phase of up to 3 seconds. In the pre-meal choice test, if participants selected a stimulus linked to an odor, the odor was delivered for 2 seconds after their choices. No odors were delivered during the post-meal choice test. Participants received the odors chosen in five randomly selected trials at the end of the task. The inter-trial interval ranged from 4 to 8 seconds, and choice and response times were recorded from all trials. Pre- and post-meal choices for both set A and set B stimuli were highly correlated (Figure S3C), indicating consistent choices across sets based on odor preferences. Thus, to assess the satiation effect on choices, we used the pre-meal average choice from set A as a session-wise odor preference baseline and compared it with the post-meal choices.

#### MRI data acquisition

MRI data were acquired on a Siemens 3T PRISMA system equipped with a 64-channel head-neck coil. Each TMS session on Day 1 and Day 2 was immediately followed by a resting-state MRI scan. Resting-state fMRI data were collected across all seven sessions with the same multi-echo sequence (310 volumes; TR = 1.5s; TE1-TE3 = 14.60ms, 39.04ms, 63.48ms). The short TE of the first echo is beneficial to mitigate signal dropout near the OFC, as demonstrated in previous studies using both resting-state and task-based fMRI.^79–82^ Other scanning parameters included: flip angle, 72°, slice thickness, 2mm (no gap), multi-band acceleration factor 4, 60 slices with inter-leaved acquisition, matrix size 104 × 104 voxels, and field of view 208mm x 208mm. A 1mm isotropic T1-weighted structural scan was acquired on Day 0 session for neuronavigation during TMS and to aid spatial normalization.

#### Coordination selection for network-targeted TMS

The stimulation coordinates were computed based on the multi-echo resting-state MRI data collected on the initial study visit. We defined our stimulation targets in the right hemisphere’s aOFC and pOFC using MNI coordinates: aOFC [34, 54, −14] and pOFC [28, 38, −16]. The pOFC coordinates were identical to those used in our previous network-targeted TMS studies.^6, 31, 49, 83^ Each targeted coordinate in the aOFC and pOFC exhibited strong functional connectivity with isolated clusters in the LPFC with peak coordinates of [44, 28, 38] and [46, 38, 14], respectively, as determined in data from Neurosynth.org involving a sample of 1,000 subjects.

We first generated spherical masks of 8-mm radius around these four coordinates in MNI space, each inclusively masked by the gray matter tissue probability map provided by SPM12 (thresholded at *>* 0.1). We transformed these four masks to each subject’s native space using the inverse deformation field generated during the normalization of the T1 anatomical image. We then specified two resting-state fMRI functional connectivity analyses (one per region) for each subject, using individual aOFC and pOFC masks as the seed regions and motion parameters from the realignment of the first echo as regressors of no interest. Stimulation coordinates were defined as the voxels within the right LPFC masks with the strongest functional connectivity to the right aOFC and pOFC seed regions, respectively. We used infrared MRI-guided stereotactic neuronavigation (LOCALITE) to apply stimulation to these two individual LPFC coordinates.

#### Transcranial magnetic stimulation

Similar to our previous work, the target coordinates were defined as the locations in the right LPFC with the strongest functional connectivity with the corresponding right OFC seed regions (see details above). The Figure-eight active/passive (A/P) coil was tilted so that the long axis was approximately perpendicular to the long axis of the middle frontal gyrus. TMS was administered at 80% of the rMT using a cTBS protocol. This protocol involved delivering bursts of three pulses at 50 Hz every 200 ms (5 Hz) for a total of 600 pulses over approximately 40 seconds. Stimulation was applied using a MagPro X100 stimulator equipped with a MagPro Cool-B65 A/P butterfly coil (MagVenture). Previous work has demonstrated that this cTBS protocol at 80% MT has inhibitory aftereffects which persist for 50–60 min over primary motor cortex.84 Whereas cTBS was delivered by positioning the active side of the A/P coil to modulate neural tissue, sham cTBS was applied with the placebo side of the A/P coil, producing similar somatosensory and auditory experiences for the participant without modulating neural tissue. Ag/AgCl tab electrodes were placed on participants’ foreheads, and weak electrical stimulation was applied in synchrony with the TMS pulses to mask somatosensory effects and enhance the perceptual similarity between cTBS and sham sessions.

Participants were informed about potential muscle twitches in the face, eyes, and jaw during simulation. To assess tolerability, two single pulses were applied over the stimulation coordinates before administering cTBS. Discomfort and perceived stimulation intensity were evaluated after each TMS session. The cTBS sessions were generally rated as more uncomfortable and intense compared to the sham sessions. On a scale from 0 (not uncomfortable at all) to 10 (extremely uncomfortable), mean discomfort ratings were 3.38 for sham and 5.8 for cTBS sessions (p < 2.2 × 10^−16^, linear mixed effects model). Similarly, on a scale from 0 (not strong at all) to 10 (extremely strong), mean intensity ratings were 3.79 for sham and 6.23 for cTBS sessions (p < 2.2 ×10^−16^, linear mixed effects model). Discomfort and intensity ratings did not differ between aOFC- or pOFC-targeted cTBS (all p > 0.6). For analyses involving cTBS effects (Day 1 or Day 2 TMS), standardized discomfort and intensity ratings were used to examine correlations or regressions against other variables, assessing if the observed cTBS effects were driven by subjective discomfort or perceived TMS intensity, but none of the effects can be explained by those ratings (see Figure S6).

### QUANTIFICATION AND STATISTICAL ANALYSIS

#### Analysis of discrimination learning

We examined whether participants improved their performance across blocks by fitting the following mixed-effects logistic regression models:

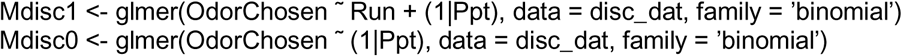

In these models, OdorChosen indicates whether the odor-predictive stimulus was selected (yes = 1) and Run ranges from 1 to 5. To assess learning across blocks, we compared a full model (Mdisc1) that included Run as a fixed effect with a reduced model (Mdisc0) that did not.

To further examine the effects of TMS and session number on discrimination learning, we grouped participants based on the session in which they received cTBS or sham stimulation on Day 1 (Figure S1B). This analysis revealed that the impairment in discrimination due to cTBS was evident only when cTBS was administered during the first session (p < 2.2 × 10^−16^). We also tested whether the effect differed by stimulation target (anterior vs. posterior OFC) but found no significant interaction or main effect related to target location (all p > 0.05).

#### Modeling discrimination learning

We used a hierarchical Bayesian implementation of the Rescorla–Wagner model^85^ to quantify value learning during the discrimination task. On each trial, participants chose between two stimuli: one predictive of an odor and one predictive of clean air. Because stimulus pairs did not repeat across trials, we modeled learning as driven primarily by the odor-predictive stimulus, tracking its value *w* over time. This value was updated based on the prediction error—the difference between the actual outcome (*w* = 1) and the expected value—scaled by a stimulus-specific learning rate α. Values were initialized at *w* = 0.5, reflecting chance-level knowledge, and progressed toward 1 with learning.

Each Day 1 session consisted of five blocks of 24 trials, covering 12 unique stimulus pairs, each presented twice per run with left/right positions counterbalanced. For a given stimulus pair, when it appeared on trial *i*, its value was updated according to:

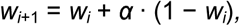

where α denotes the learning rate for that stimulus pair.

The discrimination response on trial *i*, denoted Resp*_i_*, was modeled as:

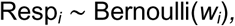

where Resp*i* = 1 if the odor-predictive stimulus was chosen, and 0 otherwise.

We estimated a separate learning rate for each odor-predictive stimulus using a hierarchical Bayesian framework with session-wise priors. Trial-level learning rates for each stimulus pair, denoted α*_j,c,p_*, were modeled as:

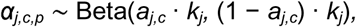

where:

- *j* indexes participants,
- *c* indexes sessions (1, 2, 3),
- *p* indexes cue pairs (1 to 12),
- *a_j,c_* is the subject- and session-specific mean learning rate,
- *k_j_* ∼ Gamma(1, 0.1) for each participant *j* = 1*, … , n*_subs_.

Higher-level learning rate means were drawn from:

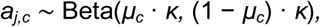

where:

- *µ_c_* ∼ Beta(8, 2) is the session-specific population mean,
- *k* ∼ Gamma(1, 0.1) controls the overall precision.

This modeling approach enabled us to derive individualized value trajectories for each odor-predictive stimulus, which were subsequently used to analyze probe choices on Day 2. Model simulations of accuracy based on posterior estimates of w closely tracked the observed data (Figure S1A), indicating that the learning model captured participants’ behavior well and was suitable for subsequent analyses.

Although not part of our original hypothesis—and rarely examined in outcome devaluation studies—we found that individual choices were also influenced by the learned value of each stimulus. The probability of choosing the SA over the NS option increased significantly with the value difference between the two stimuli (*w_SA_*− *w_NS_*), as reflected by a strong positive correlation (Pearson’s *r* = 0.92, *p* = 3.49 × 10^−10^; Figure S2).

Accordingly, when evaluating the effects of cTBS (applied on Day 1 or Day 2) on SA choices during Day 2, we included both the learned value difference (*w_SA_* − *w_NS_*) and the selective satiation index as regressors to account for factors influencing behavior beyond the effects of TMS.

#### Analysis of odor pleasantness rating

Odor pleasantness ratings were collected on a raw scale from –10 to 10. For statistical analyses, ratings were z-scored within each participant across all experimental sessions to account for individual differences in scale use. To evaluate whether selective satiation specifically reduced the pleasantness of the sated odor, we calculated the change in pleasantness (PleasantChange, defined as post-meal minus pre-meal) for each odor and session. We then fit two linear mixed-effects models with random intercepts for participants. The null model (MPC0) included only a random intercept, while the full model (MPC1) included an additional fixed effect of (IsSated), a binary variable indicating whether the odor was the sated one. Model comparison was performed using a likelihood ratio test.

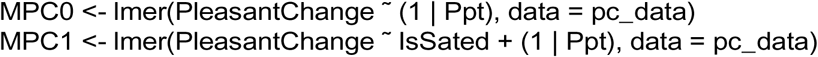

We computed a session-wise index of the selective satiation effect, SatIdx, defined as the difference in PleasantChange between sated and non-sated odors. To explore whether this effect was influenced by additional factors — such as TMS condition (Day 2; sham vs. cTBS), TMS target site (aOFC vs. pOFC), session number (1^st^, 2^nd^, 3^rd^), or sated odor type (savory/sweet) — we fit a second set of linear mixed-effects models. Each model included one of these predictors and was compared against the same null model MSatIdx0. For example, to test the influence of TMS condition (TMScond), we fit and compared the following models:

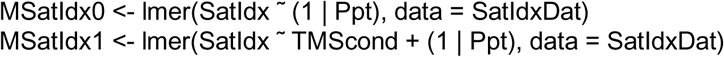

Moreover, the proportion of SA choices was significantly correlated with the pleasantness difference between sated and non-sated odors, both before and after the meal (Figure S3A), indicating that choices reflected relative odor preferences as expected. To quantify the behavioral impact of subjective value changes, we computed a “selective satiation index” by subtracting the change in pleasantness ratings for non-sated odors from those for sated odors (post-meal minus pre-meal). This index was significantly correlated with the corresponding change in SA choices (Pearson’s *r* = 0.3, *p* = 2.4 × 10^−4^; Figure S3B), further supporting a link between subjective devaluation and behavioral change.

All mixed-effects models were fit using the lme4 package in R.

To test whether the pleasantness of the two selected food odors fluctuated across sessions, we conducted a repeated-measures ANOVA on pre-meal odor pleasantness ratings across the three sessions. There was no significant main effect of session (*F* (2, 93) = 0.12*, p* = 0.887, η^2^ *<* 0.01, mean ratings: Session 1: *M* = 0.195 ± 0.080; Session 2: *M* = 0.256 ± 0.081; Session 3: *M* = 0.221 ± 0.080).

Bonferroni-corrected pairwise comparisons confirmed no significant differences between any session pairs (all *p* = 1.000), indicating that odor preference remained stable over time.

#### Analysis of probe choice responses

We analyzed the Day 2 probe data on sweet–savory choices and split these analyses by TMS target site (aOFC and pOFC groups). We used the pre-meal average choice for each session as a baseline measure of odor preference BasePref.

To assess the effect of Day 2 TMScond (Day 2; sham vs. cTBS) on choices involving the sated odor, we analyzed the trial-wise data using logistic mixed-effects modeling. The models included the following covariates: (1) BasePref, the pre-meal baseline preference; (2) SatIdx, the session-wise reduction in pleasantness of the sated odor; and (3) ValueDiff, the value difference between the two choice options on each trial, reflecting discrimination learning from Day 1. For each target group (aOFC and pOFC), we compared a full model (Mchoice1) that included the TMS condition (TMScond) with a reduced model (Mchoice0) that did not:

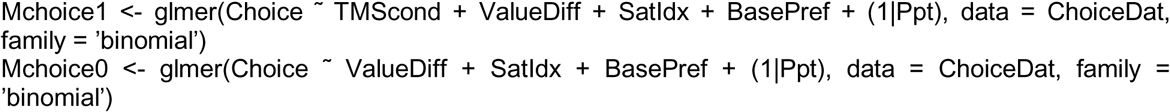

In these models, Choice was a binary outcome indicating whether the participant chose the sated odor (1) or the non-sated odor (0). To further examine whether the effect of TMS condition varied by stimulation site, we tested an additional model that included an interaction term between TMScond and TMStarget. We used the fitted function in R to extract trial-level predicted choices based on the best-fitting model for each group. These predicted values were then averaged within each participant to estimate the model-derived probability of choosing the sated odor, as shown in Figure 4. The Day 1 TMS effect was analyzed in a similar manner, using the contrast between Day 1 sham and cTBS while holding Day 2 TMS constant at sham, as shown in Figure 5.

To assess robustness, we conducted a leave-one-subject-out sensitivity analysis across key conditions and timepoints. For significant effects (pOFC Day 2 and aOFC Day 1), results remained strong and consistent (median *p* = 5.1×10^−4^ and *p* = 0.0143, respectively), with significance retained in nearly all iterations. For null effects (aOFC Day 2 and pOFC Day 1), outcomes were stable (median *p* = 0.622 and *p* = 0.263), indicating that no single participant drove the result. Even in cases where significance was not reached, effect sizes remained aligned with the group-level trend. Together, these analyses confirm that both significant and null findings were robust to the exclusion of any individual participant.

#### Multi-echo MRI data processing

Preprocessing of the multi-echo resting-state fMRI data involved several steps. First, all functional images from the smallest echo across all rs-fMRI sessions were realigned to the first volume of the first echo, and the resulting voxel-to-world mapping matrix was applied to the other two echoes, volume by volume. All functional images were then resliced for each echo. Next, the images across the three echoes were combined using temporal signal-to-noise ratio (tSNR) weighting, following parallel-acquired inhomogeneity desensitized (PAID) approach.^80^ Specifically, voxel-wise tSNR maps were computed for each echo, multiplied by the echo time (TE), and normalized across the three echoes to generate weight maps. These weight maps were then used to combine the resliced images by multiplying each volume by its respective weight map. Lastly, the combined data underwent coregistration, normalization, and smoothing using a 6 mm FWHM Gaussian kernel.

We analyzed participants’ motion during the resting-state scan after different types of TMS (sham vs. cTBS) and stimulation targeted locations (anterior vs. posterior OFC). Framewise displacement (FD) was calculated per volume and summed across volumes.^86^ No significant main effects were observed for TMS type or stimulation location (TMS type: *p* = 0.78; location: *p* = 0.94). For cTBS, FD was 39.8 mm (±13.8 mm) at the anterior OFC and 41.1 mm (±16.3 mm) at the posterior OFC; for sham, FD was 40.3 mm (±15.9 mm) at the anterior OFC and 39.5 mm (±15.7 mm) at the posterior OFC.

#### Resting-state functional connectivity analysis

Following echo combination and initial preprocessing, we further denoised the resting-state fMRI data prior to functional connectivity (FC) analysis by applying voxel-wise nuisance regression within gray matter voxels, consistent with our previous approach.^49^ The regressors included mean time series from white matter, CSF, and gray matter, motion parameters from realignment, and a linear drift term. All regressors were z-scored and included an intercept and were regressed out from the gray matter time series via linear regression. The resulting residuals were used for subsequent FC analysis.

We computed two types of FC matrices for each fMRI sessions (seven sessions total: one from the initial study visit and six from three repeated sessions on Day 1 and Day 2). First, we calculated pairwise correlations among four key ROIs—two TMS-targeted seed OFC regions and two LPFC targets—yielding a 4 × 4 FC matrix per session, used in the analyses shown in Figure 2. Second, we computed FC between the four experimental ROIs and all 116 regions from the AAL atlas^87^ for the cVAE analysis below. Due to missing signals in some atlas regions, the final FC matrix had a final dimensionality of 4 × 102.

#### Latent embedding modeling of functional connectivity of resting-state fMRI

To evaluate whether cTBS modulates brain network activity differently relative to sham, we applied a conditional variational autoencoder (cVAE) to brain functional connectivity derived from resting-state fMRI data. Our goal was to learn a low-dimensional latent representation that captures stimulation-induced variation while accounting for individual differences.

Although prior applications of VAEs and cVAEs in fMRI have largely focused on identifying individuals based on brain activity patterns,^56, 57^ our approach uses participant identity as a conditioning factor in order to isolate stimulation-induced network changes, rather than to model individual identity per se. Specifically, we conditioned both the encoder and decoder on participant identity to model subject-specific variance during reconstruction of the FC vectors. This allowed the model to account for stable inter-individual differences and focus on condition-specific effects (null, sham, or cTBS).

For each of the four ROIs, we extracted its functional connectivity (FC) pattern—defined as its correlation with all AAL regions—and used it as input to a separate cVAE model. Each model learned a latent space capturing variation in FC patterns across sessions, conditioned on individual identity. The input to the model consisted of the FC vector (*X* ∈ R^102^) and a one-hot encoded participant identity (*Y* ∈ R^48^), both of which were provided to the encoder and decoder (Figure 6). Of the 48 participants, 5 sessions were missing, resulting in a total of 331 sessions used for training.

The cVAE was implemented in PyTorch (version 2.7.0). To ensure reproducibility, all training runs were performed with a fixed random seed (42). The encoder first applies a fully connected layer (64 units, ReLU activation), followed by two linear layers that output the mean and log-variance of the latent distribution (*µ,* log σ^2^ ∈ R^10^). Latent vectors are then sampled using the reparameterization trick:

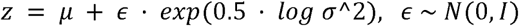

The decoder receives the concatenation of z and the condition Y, passing it through a hidden layer (ReLU, 64 units) and outputs the reconstructed FC pattern vector. The model was trained using the Adam optimizer (learning rate = 3e-4). The loss function combined mean squared error for reconstruction and the KL divergence between the approximate posterior and the standard Gaussian prior:

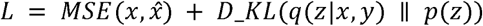

To quantify TMS effects, we calculated the Euclidean distance between each session’s latent embedding (for cTBS or sham) and that of the null condition baseline. This distance served as a proxy for the degree of deviation from unstimulated resting-state activity. Because the latent space is optimized to capture meaningful variation in functional connectivity while controlling for individual differences, larger distances were interpreted as reflecting stronger TMS-induced modulation.

Prior to model training, each FC feature was z-scored to zero mean and unit variance across sessions (sham, cTBS, null). Because global standardization does not force equality of variances or means across groups, we additionally tested both variance and magnitude across TMS conditions. For variance, Brown–Forsythe tests with Benjamini–Hochberg FDR across 102 features found no differences (0/102 at q < 0.05); median within-group variances were similar (cTBS = 0.984; null = 1.003; sham = 1.012), and a pooled test showed no effect (F = 0.347, p = 0.707). For magnitude, Welch’s one-way ANOVAs with FDR likewise detected no feature-level mean differences (0/102 at q < 0.05); median group means were near zero (cTBS = −0.0068; null = 0.0060; sham = −0.0022), and a pooled test of subject-level averages confirmed no differences (F = 0.019, p = 0.981, η² ≈ 0). Thus, the standardized inputs were well matched across conditions in both variance and mean.

To ensure that model performance was comparable across conditions, we assessed reconstruction fidelity after model training by applying the trained cVAE to all sessions and computing reconstruction accuracy separately for each session. For each session, we calculated the Pearson correlation between the original and reconstructed functional connectivity matrices. We chose correlation as our primary measure because it is scale-invariant, capturing the preservation of the connectivity pattern rather than absolute differences in scale. Correlation values were nearly identical for sham (*r* = 0.8463) and cTBS (*r* = 0.8466) conditions (*t* = −0.028*, p* = 0.978), indicating that the decoder’s performance was comparable across conditions. These results suggest that observed differences in latent distances are unlikely to be driven by differences in reconstruction accuracy.

One important consideration in our analysis was the inherent imbalance between sham and cTBS sessions due to the experimental design. For participants without missing data, each contributed two cTBS sessions and four sham sessions. As a result, the cVAE may have become better optimized to the FC patterns of sham sessions, since all sessions were used simultaneously during training. To address this potential bias, we repeated the cVAE training for each ROI using a resampling strategy. Specifically, during each training epoch, we applied weighted random sampling of sessions, assigning weights inversely proportional to the number of sessions per condition. This ensured a more balanced representation of sham and cTBS conditions throughout training. The repeated modeling yielded results consistent with those from the original model, suggesting that our findings were not driven by session imbalance.

To evaluate whether our findings depended on the choice of latent dimensionality (i.e., 10), we conducted a sensitivity analysis by retraining the cVAE with a range of latent dimensions of 2, 4, 8, 12, 16, and 32, while keeping all other model parameters and training procedures constant. For each model, we computed Euclidean distances in the latent space between each of the sham and cTBS conditions and the null condition and compared sham versus cTBS distances using paired *t*-tests across subjects (*n* = 46). We observed significantly greater distances in the cTBS than the sham condition relative to the null condition for latent dimensions of 8 (*t* = −2.35, *p* = 0.023), 12 (*t* = −2.47, *p* = 0.018), 16 (*t* = −2.29, *p* = 0.027), and 32 (*t* = −2.08, *p* = 0.043). However, we found no significant effects for 2 (*t* = −0.53, *p* = 0.60) and 4 (*t* =−1.35, *p* = 0.18) latent dimensions. The absence of significant effects at 2 and 4 latent dimensions may reflect insufficient representational capacity in the latent space. When the dimensionality is too low, the model likely prioritizes capturing the largest sources of variance (e.g., subject-specific structure) at the expense of finer, condition-related differences, leading to reduced sensitivity in the Euclidean distance measure. These results indicate that the observed effect is robust across a range of moderate-to-high latent dimensionalities (≥ 8) but may require a minimal representational capacity to detect stimulation-induced changes.

## SUPPLEMENTAL INFORMATION

Document S1. Figures S1–S6

