## Supplemental Information for "Distinct contributions of anterior and posterior orbitofrontal cortex to outcome-guided behavior"

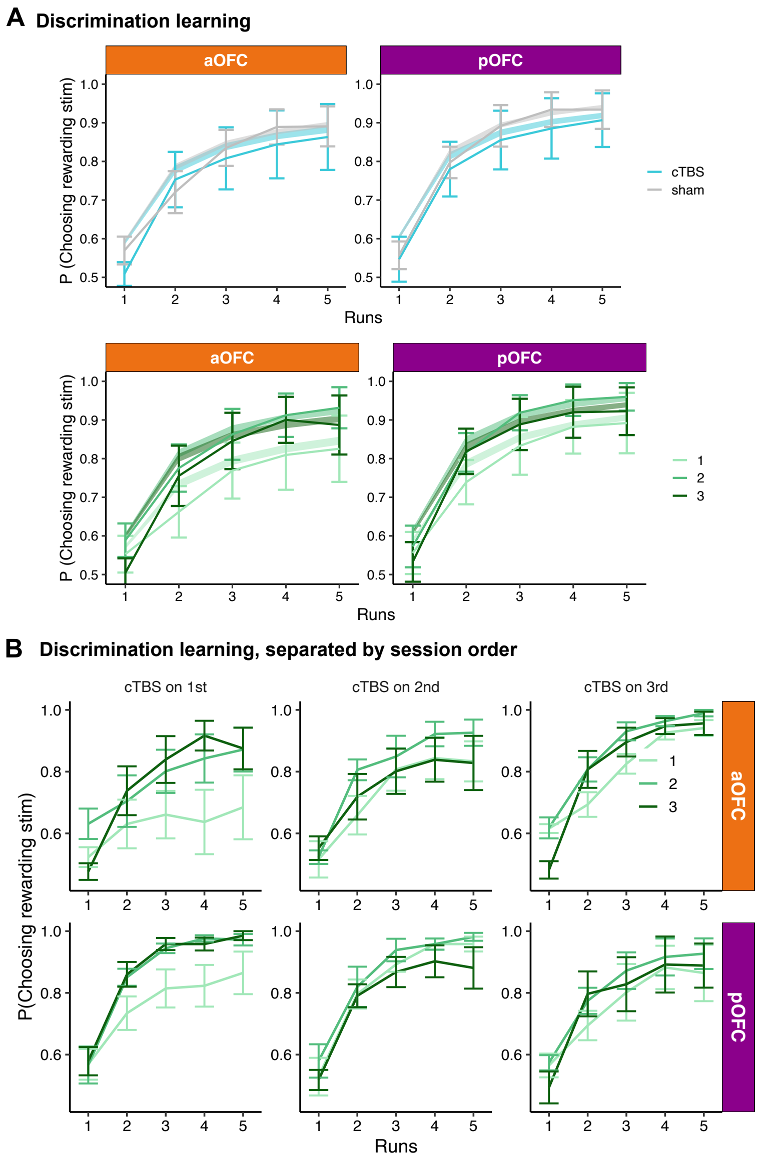


Figure S1: **Posterior or anterior OFC-targeted cTBS disrupted value acquisition during the first session, related to Figure 3. A.** Discrimination accuracy over five runs during the Day 1 task, plotted by TMS condition (cTBS vs. sham), session number ($1^{st}$, $2^{nd}$, $3^{rd}$), and stimulation target (aOFC vs. pOFC). Line plots with error bars represent observed data (mean $\pm$ SEM), while shaded regions indicate the 95% confidence intervals simulated from posterior estimates of stimulus weights ($w$, see Methods). **B.** Discrimination accuracy across runs, separated by session number and the session order of cTBS administration—that is, whether cTBS was applied during the $1^{st}$, $2^{nd}$, or $3^{rd}$ session of the three-session experiment.

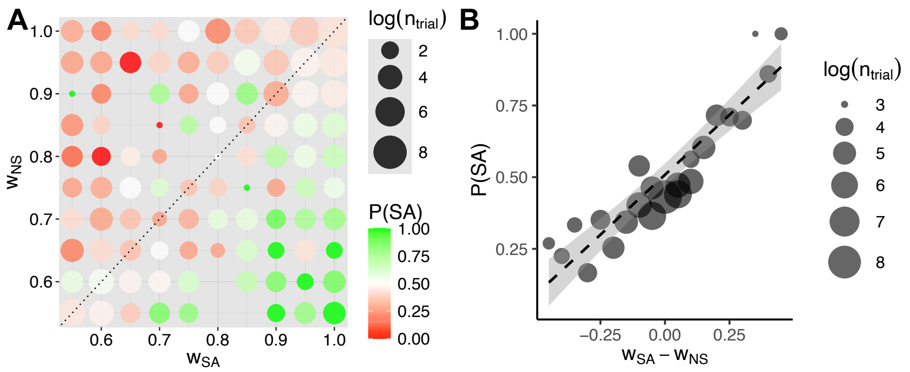


Figure S2: **Probe choices are influenced by learned stimulus values, related to Figures 4 and 5. A.** Choice of sated odors options associated with each of the learned weight of the combination of sated and non-sated options. Dot size represents the number of trials per value combination (log-scaled), with missing dots indicating unobserved combinations. **B.** Probability of choosing the sated odor stimulus as a function of the estimated value difference between the sated and non-sated options ($w_{SA} - w_{NS}$). Dot size reflects the number of trials (log-scaled) at each value bin.

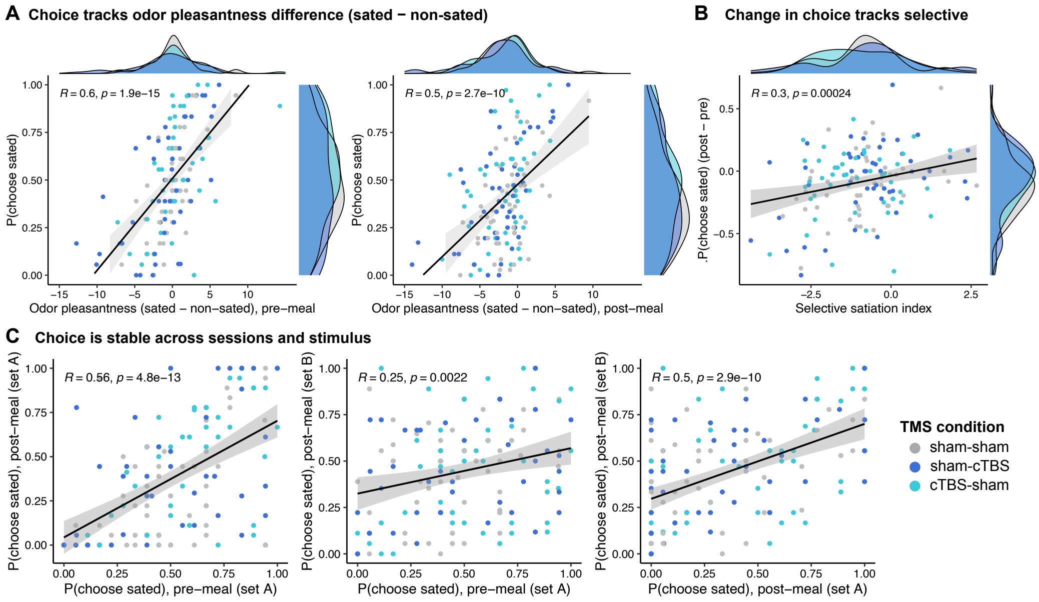


Figure S3: **Odor pleasantness ratings and selective satiation predict sated-odor choices; preferences are stable across sessions and stimulus sets, related to Figure 3, 4, 5. A.** Across participants, the probability of choosing the sated-odor cue increases with the pleasantness difference between sated and non-sated odors both pre-meal (left) and post-meal (right). **B.** The change in choice from pre to post meal ($\Delta P_{choose-sated}$) is positively related to the selective satiation index (change in pleasantness for the sated odor relative to the non-sated odor from pre to post). **C.** Scatter plots showing correlations in the proportion of sated odor choices across sessions (pre- and post-meal) and stimulus sets (A and B).

Dots are participants (color = TMS condition); black lines show least-squares fits with 95% CI; ridge plots show marginal distributions. Pearson's r and p values are computed across all participants (collapsed across TMS conditions).

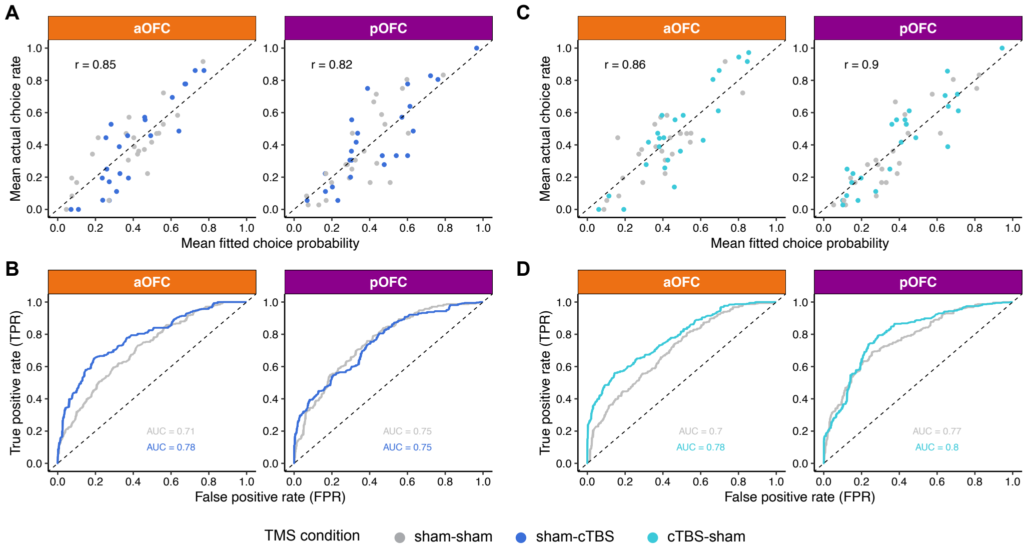


Figure S4: **Model fit evaluation across stimulation sites and conditions for Day 2 and Day 1 TMS effect, related to Figures 4 and 5.** **A.** Across-participant correlation between the mean fitted choice probability and the actual mean choice rate for each subject, shown separately for aOFC (left) and pOFC (right) groups. Each dot represents a single subject, colored by condition. The dashed diagonal line indicates perfect correspondence between model predictions and behavior. **B.** Receiver operating characteristic (ROC) curves for predicting trial-level choices from model-estimated probabilities, shown separately for each stimulation site and condition. The area under the curve (AUC) indicates the model's discriminative ability; higher curves reflect better separation between choice = 1 and choice = 0. Dashed line represents chance-level performance. **C–D.** Same analyses as in A and B, but for the Day 1 TMS effect. *Note: The ROC curves reflect trial-level discriminability of the model predictions, whereas Figures 4 and 5 capture overall choice bias. Therefore, differences between conditions observed here may not necessarily appear in the other measure, and vice versa.*

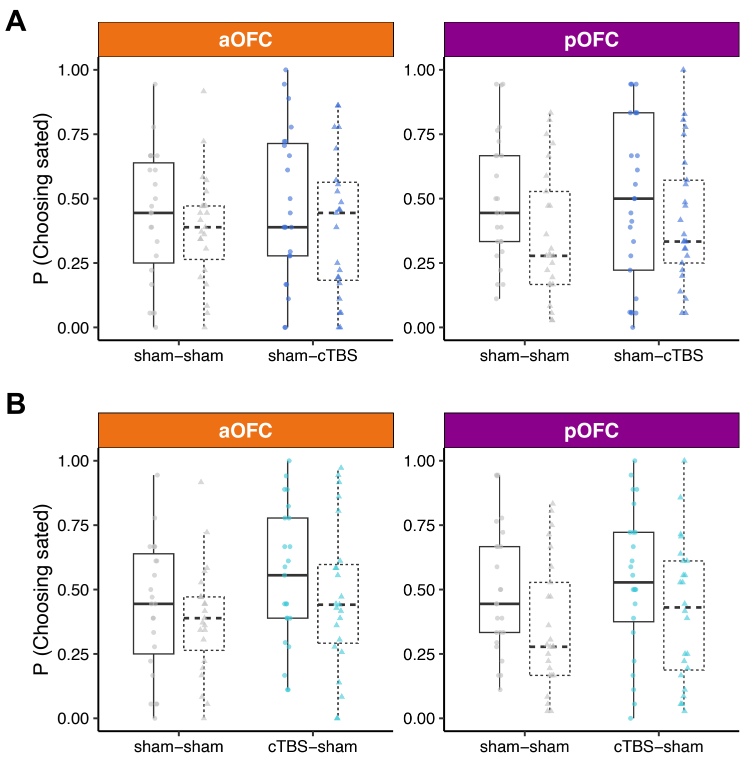


Figure S5: **Raw data for Day 2 and Day 1 TMS conditions on sated odor choices, related to Figures 4 and 5. A.** Choice behavior for sated odors before and after the meal, shown separately by stimulation site (aOFC vs. pOFC) and stimulation condition (sham–sham vs. cTBS–sham). These data correspond to the Day 2 TMS manipulation and provide the basis from which the Day 2 effect is derived (see Figure 4). Boxplots display the distribution of choice probabilities across participants, with lines connecting Pre- and Post-meal values for the same participant within each condition. Dots indicate Pre-meal choices; triangles indicate Post-meal choices. A reduction from Pre to Post reflects a successful selective devaluation effect. **B.** Same format as A but for sham–sham and sham–cTBS conditions on Day 1, providing the data used to derive the Day 1 TMS effect (see Figure 5).

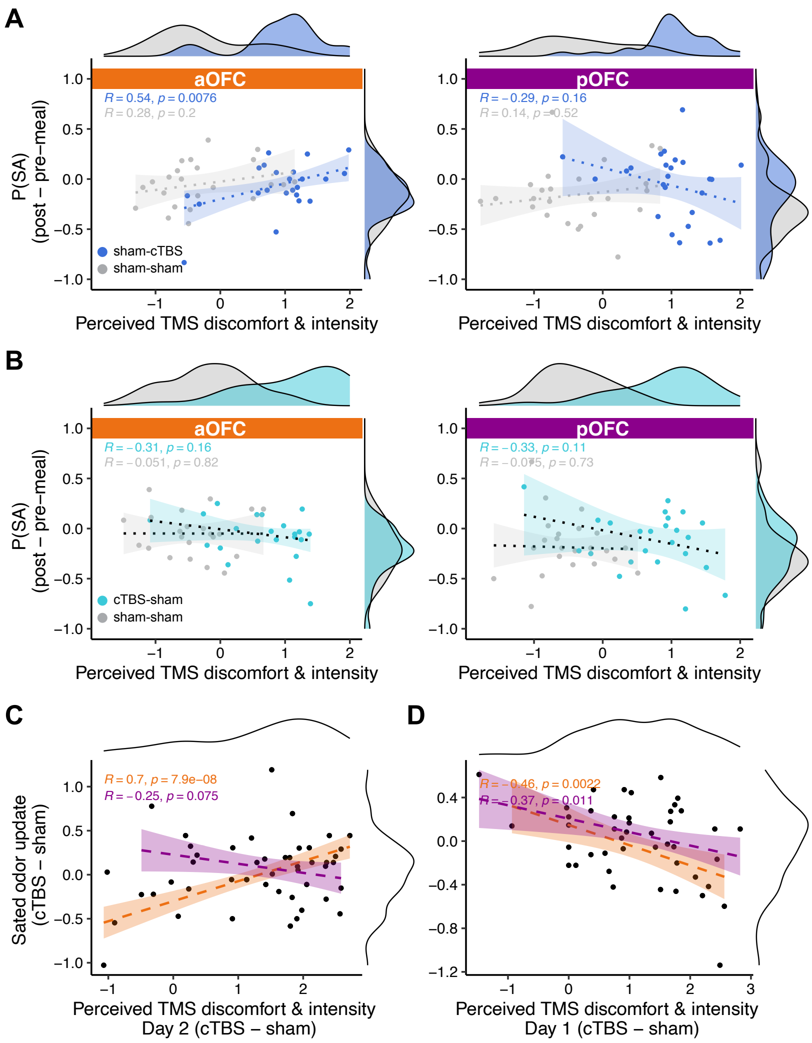


Figure S6: **Relationship between perceived TMS discomfort and intensity and sated odor (SA) choices, related to Figures 4 and 5. A.** Correlation between SA choices and TMS ratings, separated by Day 2 TMS conditions (sham-cTBS vs. sham-sham) and TMS targeted regions (aOFC, pOFC). A positive correlation was observed between TMS ratings and SA choices in the aOFC group, but including ratings of TMS perception into the regression models did not alter the observed TMS effects on SA choices. **B.** Same as **A**, but focus on Day 1 TMS effect (sham-sham vs. cTBS-sham). **C.** Scatter plot showing the relationship between the condition-wise difference in SA choices (sham–cTBS minus sham–sham) and the corresponding difference in TMS intensity ratings on Day 2. A significant positive correlation was observed in the aOFC group ($p = 7.9 \times 10^{-8}$). **D.** Same as **C**, but focus on Day 1 TMS effect (sham-sham vs. cTBS-sham). Shaded areas represent 95% confidence intervals estimated using robust linear regression. Marginal distributions are shown on the top and right axes.
